# Capturing transient states of heterodimeric ABC transporter TM287/288 by Time-Resolved Small-Angle X-ray Scattering

**DOI:** 10.1101/2025.09.29.679171

**Authors:** Lea Schröder, Dario De Vecchis, Andrey Gruzinov, Lars V. Schäfer, Clement E. Blanchet, Markus A. Seeger, Henning Tidow, Inokentijs Josts

## Abstract

Structures of the heterodimeric ABC transporter TM287/288 have been previously determined in several states indicating large conformational changes during its reaction cycle. However, for a complete description of the cycle, transient states (such as an occluded state) are still missing, as they are difficult to capture with static structural biology techniques. Here, we investigate the kinetics and accompanying structural changes in the course of the reaction cycle of full-length TM287/288 using time-resolved small-angle X-ray scattering, initiated by stopped-flow mixing. The use of active site mutants and state-specific sybodies/nanobodies enabled us to dissect the temporal events involved in the ATP-driven conformational cycle of TM287/288 and reveal a transient occluded state of this ABC transporter.

**Statement of Significance:** ATP-binding cassette transporters are essential molecular machines that use the power of ATP binding and hydrolysis to move substances across cellular membranes. Here, we use time-resolved small-angle X-ray scattering combined with conformation-specific nanobodies to directly capture and assign different states to the heterodimeric ABC transporter TM287/288 in solution. Our findings advance our understanding of ABC transporter and demonstrate a broadly applicable method for dissecting the different molecular states in membrane proteins on a second-to-minute time scale.

**TOC figure:** 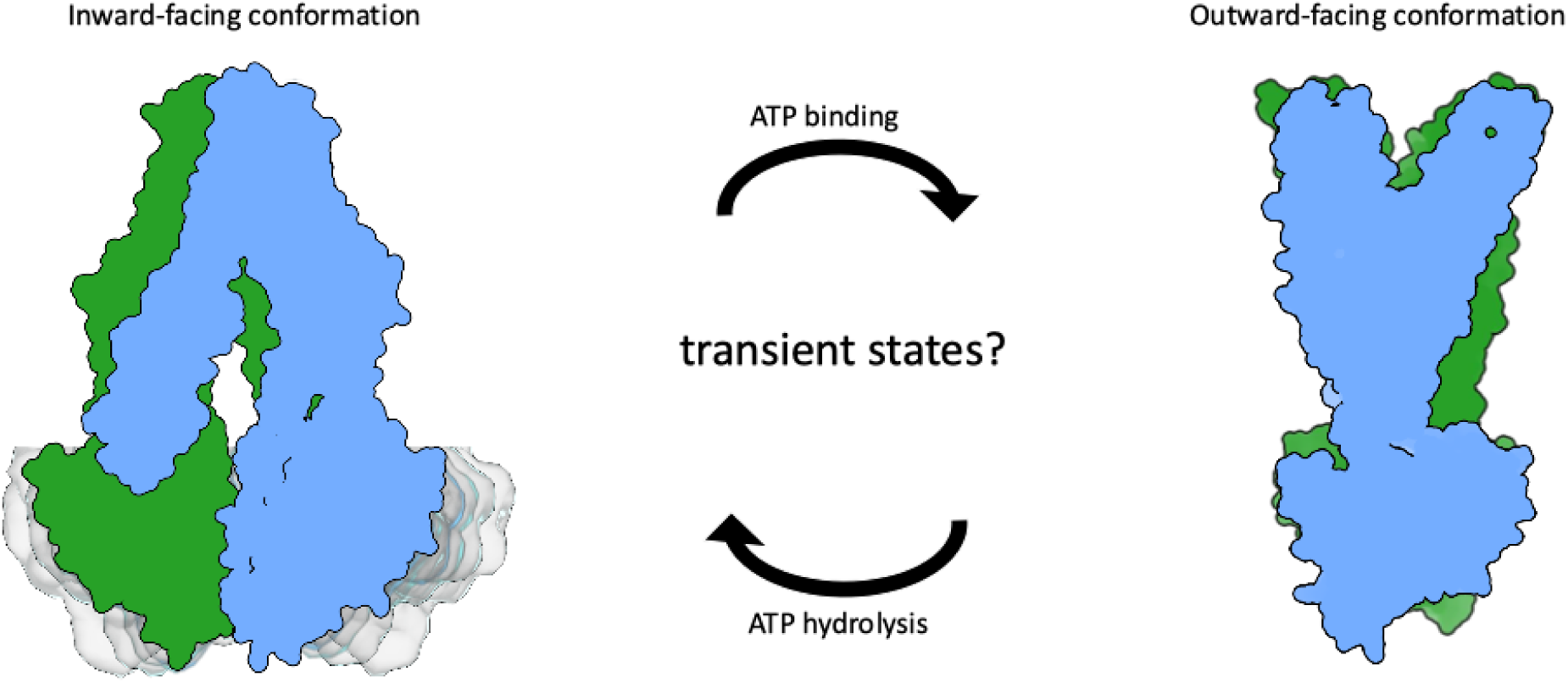

## Introduction

ATP-binding cassette (ABC) transporters are ubiquitous integral membrane proteins found across all domains of life [1–3]. These transporters mediate the ATP-dependent translocation of a wide range of substrates—including ions, lipids, peptides, and large organic molecules—across biological membranes [4–7]. Based on their transport directionality, ABC transporters are generally classified as importers, exporters, or extruders [8]. They operate as homo- or heterodimers, coupling ATP-Mg²⁺ binding, ATP hydrolysis, and ADP/Pi release to substrate translocation via a mechanochemical mechanism [9].

ABC transporters share a conserved architecture composed of two transmembrane domains (TMDs) and two nucleotide-binding domains (NBDs). The TMDs define substrate specificity and form the translocation pathway, whereas the NBDs supply the free energy required for transport via ATP binding and hydrolysis. Early mechanistic descriptions framed this coupling in terms of a nucleotide-driven conformational “power stroke,” in which ATP-induced NBD dimerization promotes large-scale rearrangements of the TMDs [9–11]. Subsequent biochemical, structural, and spectroscopic studies have demonstrated that this process is better described as a series of coordinated and reversible conformational equilibria linking nucleotide occupancy at the NBDs to alternating-access transitions of the TMDs, rather than a single discrete mechanical step [12–14].

In ABC exporters, the transport cycle is commonly described by an alternating-access mechanism in which the resting inward-facing (IF) conformation with separated NBDs transitions to an outward-facing (OF) conformation upon ATP-induced NBD dimerization, followed by ATP hydrolysis, NBD disengagement, and return to the IF state [14]. Although still debated, ATP binding alone seems to be sufficient for the IF-to-OF transition while ATP hydrolysis and phosphate release will subsequently reset the transporter to the IF state [15–17].

Numerous structures of ABC transporters have been determined in various conformational states (IF/Occ/OF) providing distinct snapshots of the conformational cycle [8, 18–22]. In type IV ABC transporters communication between NBDs and TMDs is mediated via large intracellular domains containing ‘coupling helices’ [3]. Recent integrative structural, biochemical, and computational studies have further refined this model, providing detailed insight into the temporal order of nucleotide binding, hydrolysis, and conformational transitions in type IV ABC transporters [15, 17, 23, 24].

Importantly, not all ABC transporters possess two catalytically equivalent ATPase sites. Many heterodimeric ABC transporters exhibit intrinsic asymmetry, with one consensus site capable of ATP hydrolysis and one degenerate site that binds but does not hydrolyze ATP [8]. TM287/288, a heterodimeric ABC exporter from *Thermotoga maritima*, exemplifies this class. It contains one catalytically competent ATP-binding site and one degenerate site, resulting in asymmetric nucleotide processing during the transport cycle [25]. Although its native substrate is unknown TM287/288 has been shown to transport the anticancer drug daunomycin as well as Hoechst 33342 [26]. TM287/288 has been structurally characterized in multiple nucleotide-and conformation-dependent states, making it a well-established model for dissecting the mechanistic consequences of ATPase asymmetry in ABC exporters [26–28].

Recently, two conformation-specific single-domain antibodies were developed that selectively recognize and stabilize the OF conformation of TM287/288, thereby inhibiting its transport activity [28]. Sybody Sb#35 binds to the extracellular wing region of the transporter, whereas nanobody Nb#1 targets the closed NBD dimer interface from the cytosolic side, with an epitope spanning both TM287 and TM288. Binding of both antibodies is ATP dependent and results in inhibition of ATPase activity, providing powerful tools for selectively trapping defined functional states along the transport cycle.

Time-resolved small-angle X-ray scattering with stopped-flow mixing (SF-TR-SAXS) is a solution-based technique capable of capturing protein conformational changes on the millisecond timescale [29]. SF-TR-SAXS has been successfully applied to study protein folding reactions and ligand-induced quaternary structural transitions [30–33]. We previously employed SF-TR-SAXS to resolve the kinetics of NBD dimerization and dissociation in both isolated NBDs and full-length MsbA reconstituted in lipid nanodiscs, demonstrating its suitability for dissecting transporter conformational dynamics in real time [34].

In the present study, we use SF-TR-SAXS to investigate the conformational dynamics of the heterodimeric ABC exporter TM287/288. By exploiting conformation-specific single-domain antibodies to selectively stabilize distinct functional states, we assign time-resolved SAXS signatures to discrete steps of the transport cycle and identify transient intermediates that are not readily accessible by static structural methods.

Collectively, our data provide a kinetic and structural dissection of the conformational transitions underlying the functional cycle of the heterodimeric ABC transporter TM287/288, placing these observations within the context of previously established mechanistic models for type IV ABC exporters and extending them through direct experimental characterization of transient intermediates.

## Materials and Methods

### Expression and purification of full-length TM287/288

The plasmid encoding for the heterodimeric ABC transporter TM287/288 of *T. maritima* has been previously described [26]. The expression vector is pBXNH3L. Expression and purification were adapted from Hutter *et al*. [28]: Transformed *E. coli* MC1061 cells were grown in TB medium supplemented with 100 µg/mL ampicillin at 37 °C for 1.5 h and further grown at 30 °C until an OD_600_ of 1.5 was reached. Expression was then induced with 0.0017 % (w/v) L-arabinose. After 5 h cells were harvested. For membrane preparation, cells were disrupted in lysis buffer (20 mM Tris, 200 mM NaCl and 10 % (v/v) glycerol, pH 7.5) supplemented with lysozyme, DNase and protease inhibitor. For solubilization, the membrane pellet was then resuspended in lysis buffer and supplemented with 1 % (w/v) DDM. Membranes were solubilized for 2 h at 4 °C. After addition of 30 mM imidazole, the supernatant was applied onto a Ni-NTA column. The column was washed with 50 mM imidazole, 200 mM NaCl, 10 % glycerol and 0.03 % DDM at pH 7.5. TM287/288 eluted with 200 mM imidazole, 200 mM NaCl, 10 % glycerol and 0.03 % DDM at pH 7.5. The buffer was exchanged to 20 mM Tris, 150 mM NaCl and 0.03 % DDM (pH 7.5) and size exclusion was performed using a Superose 6 Increase 10/300 column.

Compared to its various pdb structures the TM287/288 protein contains an additional N-terminal His-tag and linker sequence: MHHHHHHHHHHLEVLFQGPSGSGGGGGS.

### Expression and purification of nanobody Nb#1 and sybody Sb#35

The plasmids encoding for nanobody Nb#1 and sybody Sb#35 have been previously described [28]. The expression vector is pSBinit. Transformed *E. col*i MC1061 cells were grown in TB medium supplemented with 25 µg/mL chloramphenicol at 37 °C and 100 rpm for 2 h and further grown at 25 °C until an OD_600_ of 1.5 was reached. Expression was then induced with 0.02 % (w/v) L-arabinose. After incubation overnight cells were harvested. Cells were disrupted in lysis buffer (20 mM Tris, 200 mM NaCl and 10 % (v/v) glycerol, pH 7.5) supplemented with lysozyme, DNase and protease inhibitor and afterwards centrifuged. The supernatant was supplemented with 30 mM imidazole and applied onto a Ni-NTA column. The column was washed with 50 mM imidazole, 200 mM NaCl and 10 % glycerol at pH 7.5 and Nb1/Sb35 finally eluted with 200 mM imidazole, 200 mM NaCl, 10 % glycerol and 0.03 % DDM at pH 7.5. The buffer was exchanged to 20 mM Tris and 150 mM NaCl (pH 7.5) and size exclusion performed using a HiLoad 16/600 Superdex 75 column.

Compared to the pdb model 6qv1 the Nb#1 protein contains the additional residues GRAGEQKLISEEDLNSAVDHHHHHH. Compared to the pdb models 6quz and 6qv0 the Sb#35 protein contains the additional residues AGRAGEQKLISEEDLNSAVDHHHHHH.

### (Stopped-flow time-resolved) Small-angle X-ray scattering experiments

All synchrotron SAXS data were collected at beamline P12 operated by EMBL Hamburg at the PETRA III storage ring (DESY, Hamburg, Germany) [35]. For batch measurements, TM287/288 was dialyzed against TBS buffer (20 mM Tris, 150 mM NaCl, pH 7.5) + 0.03 % DDM. TM287/288 was measured at a final concentration of 20 µM. Nb#1 in TBS + 0.03% DDM was added in either 2x molar excess (40 µM) or 10x molar excess (200 µM). All batch measurements were performed at 20 °C, with data collected on the Pilatus 6M detector. A total of 10 frames with 100 ms exposure per frame were collected for all the protein samples, with buffer frames collected before and after each protein measurement. Data were averaged and background subtracted against the appropriate buffers using automatic procedures on the beamline.

The following experiments were performed in 20 mM HEPES, pH7.6, 200 mM NaCl, 5 mM MgCl_2_. For the stopped-flow time-resolved SAXS experiments, TM287/288 purified in DDM (approx. 20 μM) was mixed with Mg^2+^-ATP (1 mM) using a stopped-flow device, simultaneously injecting 60 μl of each component with a flow-rate of 2 ml/s resulting in a dead-time of 5.0 ms (Bio-logic, Seyssinet-Pariset, France). Nb#1 and Sb#35 were loaded with TM287/288 in 2x excess in one of the syringes for the kinetic experiments. All stopped-flow measurements were performed at room temperature, with data collected on the Eiger 4M detector with a sample to detector distance of 3m. The SAXS experiments were carried out with a wavelength of 1.24 Å (10keV), transmission was set to 30% (beam flux ≈ 2.10^12^ photons/s) and X-ray exposure time was set to 25 ms. 40 frames were acquired per injection. Kinetics were captured by adjusting the delay between the sample mixing and the acquisition of the scattering data. The 40 frames collected for each injections were compared using CorMap [36] (Franke, Jeffries et al. 2015) and statistically similar frames were averaged.

SAXS data were analyzed using ATSAS [37]. R_g_ values were calculated using Primus with errors obtained from the Guinier fit.

### Modelling of TM287/288 occluded states structures

TM287/288 structural models were built with Modeller [38] using available templates. Residues 2–569 for TM287 and residues 22–591 for TM288 were considered. Nucleotides were not modeled. For the inward-facing state, pdb: 4Q4A; Wide inward-facing state, pdb: 6BL6; (Outward-facing) occluded state, pdb: 4S0F, 7PR1 and 6RAI; Outward-open state, pdb: 6QUZ; Outward-open state with Nb#1/Sb#35-bound, pdb: 6QUZ.

### Modelling and data analysis

Models of DDM detergent-micelle-embedded TM287/288 in different conformations were built using CHARMM-GUI micelle builder with 170 β-DDM molecules [39], and CRYSOL [40] was used to calculate the R_g_ of the models and theoretical scattering curves.

### Activity assays

TM287/288 activity was measured performing the Baginski assay [41, 42] as previously described [43]. Five micrograms of TM287/288 in activity assay buffer (20 mM Tris, 150 mM NaCl, 5 mM MgCl_2_, 0.03% DDM) were incubated with 0/0.05/0.1/0.2/0.35/0.5/0.75/1/2/4 mM ATP in a total volume of 50 µL for 20 min at RT (final TM287/288 conc.: 0.75 μM). Addition of 50 µL freshly prepared ascorbic acid solution (140 mM ascorbic acid, 0.5 M HCl, 0.1 % SDS, 5 mM ammonium heptamolybdate) stopped the reaction. After 10 min incubation at RT 75 mL sodium citrate solution (2 % sodium citrate, 2 % sodium metaarsenite, 2 % acetic acid) were added to stabilize the color. After approx. 45 min absorbance at 860 nm was read using a Tecan Spark 20 M. To assess how Nb#1 and Sb#35 affect activity, they were added to wt TM287/288 in 2x and 10x molar excess analogous to the time-resolved SAXS experiments. The activity assays were then performed as described above. All experiments were performed as triplicates and, for comparability to time-resolved SAXS, at RT.

### Bio-layer interferometry (BLI)

TM287/288 (wt) was biotinylated for Biolayer Interferometry (BLI) experiments. The biotinylation reaction was performed mixing TM287/288 wt in 137 mM NaCl, 2.7 mM KCl, 10 mM Na_2_HPO_4_, 1.8 mM KH_2_PO_4_ and 0.03 % DDM pH 6.5 with EZ-link sulfo-NHS-LC-LC-biotin in DMSO in a 1:5 molar ratio. The mixture was incubated at 4 °C for 24 h. Excess biotin was removed and the buffer exchanged to 20 mM Tris, 150 mM NaCl and 0.03 % DDM pH 7.5. BLI measurements were performed at 25 °C using the Octet RED96 system by Sartorius. For all steps buffer containing 20 mM Tris, 150 mM NaCl, 5 mM MgCl_2_, 1 mM ATP and 0.03% DDM (pH 7.5) was used. Streptavidin sensors were pre-equilibrated in buffer for approx. 30 min. A buffer baseline was measured for 60 s followed by immobilization of 3 µg/mL TM287/288 wt (0 µg/mL for reference) onto the sensors for 240 s. After a second baseline (120 s) the association of 40 µM Nb#1 was recorded for 120 s followed by a second association phase of 120 s with 40 µM Nb#1 and 40 µM Sb#35. Buffer was used to dissociate the analytes for 500 s.

### Data and software availability

Representative SAXS data have been deposited at the SASBDB (https://www.sasbdb.org) and have been assigned the following accession code: XXXXXX. Data for all time points are available for download from: https://www.sasbdb.org/data/xxxxxxx/. All SAXS data processing and analysis were performed using the ATSAS suite [37]. Visualization and image preparation were performed with PyMOL, Excel, Adobe Illustrator, GraphPad Prism and PowerPoint.

## Results and Discussion

### Transitions between different conformational states of TM287/288 can be investigated by SF-TR-SAXS

Using the stopped-flow setup installed at the P12 Bio-SAXS beamline at EMBL, Hamburg [35] we mixed apo-TM287/288 purified in DDM (20 μM) with Mg^2+^-ATP (1 mM) and recorded SAXS profiles at room temperature in regular intervals (from 0 s to 4 min) (Fig. 1). Mixing of Mg^2+^-ATP with purified TM287/288 led to a decrease in the radius of gyration (R_g_) of TM287/288 after 1 s, reaching a minimum at approx. 20 s (Fig. 2A). After 30 s the R_g_ increases slightly and plateaus until our latest timepoint of 240 s without returning back to the original size. Kratky plots of representative experimental SAXS curves at selected time points are shown in Suppl. Fig. S1. Scattering difference curves show that conformational changes occur in the q region 0.015-0.05 Å^-1^ implying large-scale structural changes in TM287/288 upon ATP binding (Suppl. Fig. S2). Our previous work with the ABC exporter MsbA in nanodiscs has concluded that the initial reduction in protein size can be attributed to the dimerization of the NBDs and the formation of an occluded transporter state [34]. Therefore, we interpret the phase of initial R_g_ compaction in TM287/288 as ATP-induced dimerization of the nucleotide-binding domains (NBDs) leading to an occluded state after 20-30 s. Such an occluded state of TM287/288 has not been experimentally shown before, but simulations of the catalytic cycle of TM287/288 showed that this intermediate does form *in silico* [44, 45]. Therefore, we modeled this state using PCAT1 (pdb:4S0F, [20]), CtAtm1 (pdb:7PR1, [19]) and TmrAB (pdb:6RAI, [24]) ABC transporters in (outward-facing) occluded conformation as templates (Fig. 2 / Suppl. Fig. S3). The observed change in R_g_ (ΔR_g_) in our SAXS data is consistent with the calculated R_g_ change based on structural models of TM287/288 in the different conformational states (Fig. 2B). Additionally, comparison of calculated states of TM287/288 (Occ and OF) against our experimental scattering difference curves at 25 s and 240 s support these findings (Suppl. Fig. S4 C-D).

**Figure 1.**
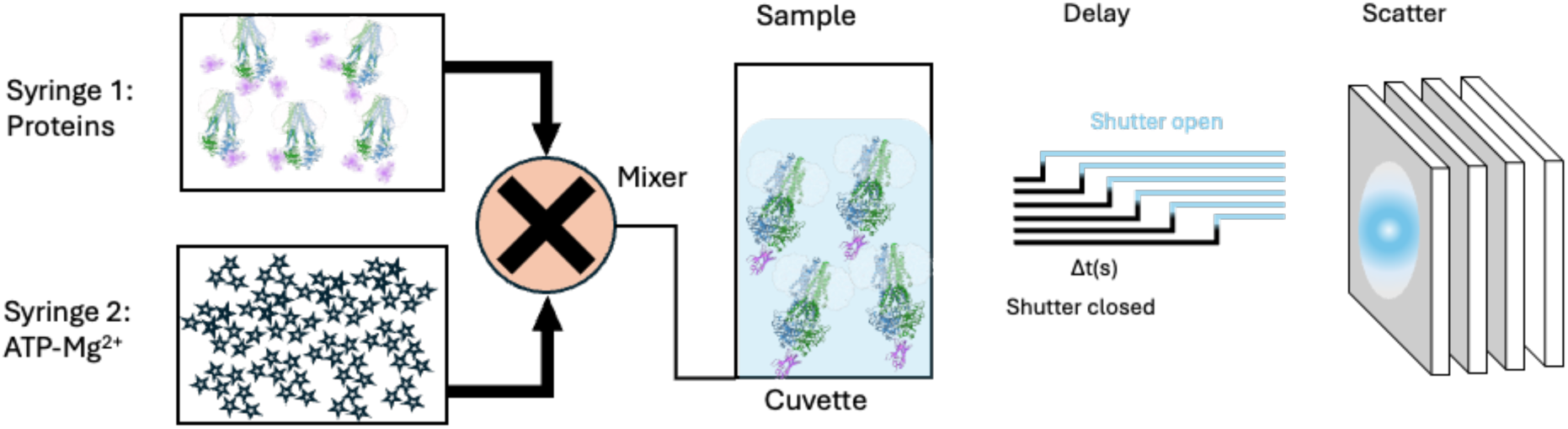
Schematic setup for time-resolved SAXS measurements with TM287/288 (and single-domain antibodies (Nb#1/Sb#35)). TM287/288 (in absence or presence of Nb#1/Sb#35) in syringe 1 was mixed with ATP-Mg^2+^ in syringe 2 and SAXS data were acquired after various time delays ranging from 0 ms to 240 s. The first measurement (zero delay) consists of the acquisition of 40 frames with 25 ms exposure each. Therefore, the first timepoint in our experiments is an average of all frames between 0 and 1 s. Subsequent delays are also averages of 1000 ms frames from the time of shutter opening.

**Figure 2.**
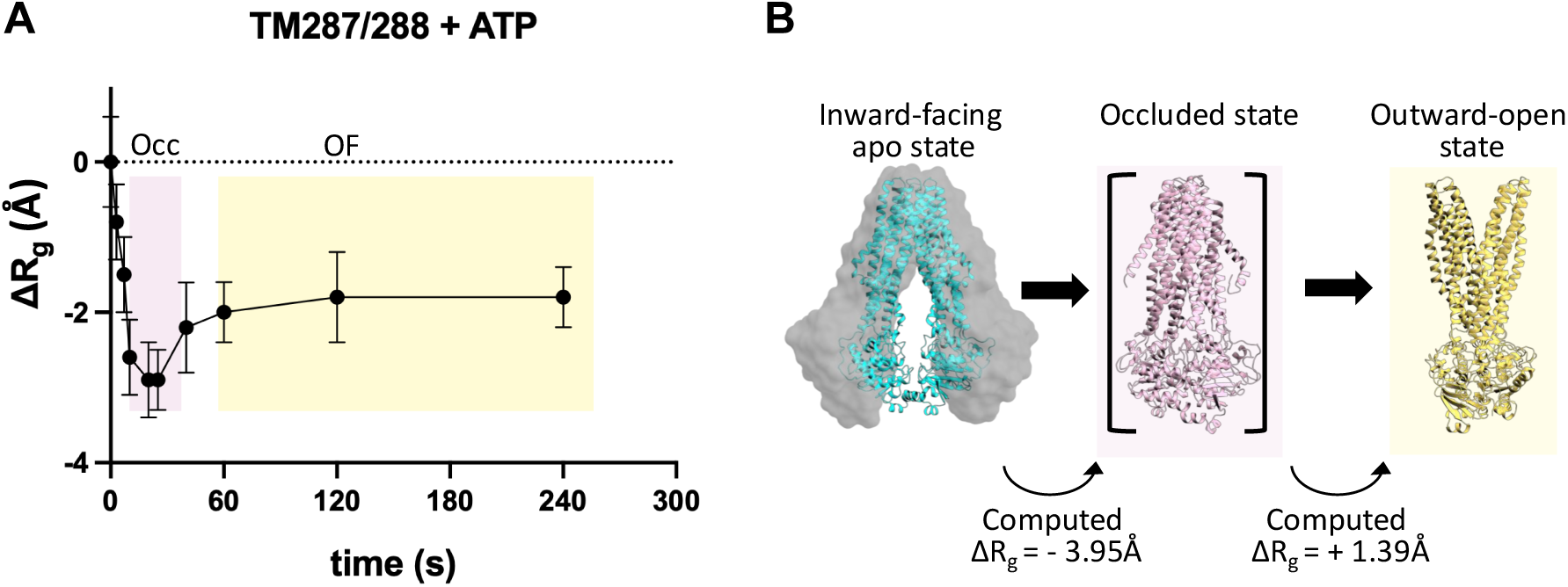
ATP-driven conformational changes of TM287/288 can be followed by TR-SAXS. A) SAXS data acquired after mixing and various delays were used to calculate changes in radius of gyration (R_g_) with errors obtained from the Guinier fit. The dashed line corresponds to the starting state. B) Conformational changes of TM287/288 models that were assigned to the phases in the TR-SAXS profile. The grey envelope corresponds to a wide-open apo model. R_g_ values were calculated with CRYSOL.

It should be noted that the assignment of the occluded state (as well as other conformational states) is based on agreement between experimental and calculated R_g_ changes, making use of existing structures in different conformational states. While this indirect assignment does not rule out alternative conformations leading to similar R_g_ changes, our assignment is supported by a wealth of prior ABC transporter studies showing that ATP-binding to the apo proteins leads to NBD dimerization and thus compaction of the transporter.

To investigate whether the increase in R_g_ after 20 s could be a consequence of ATP-hydrolysis and subsequent dissociation of the nucleotide-binding domains, we used a catalytically-inactive E/Q-mutant (TM288_E517Q_) as comparison. This E/Q-mutant has been shown not to hydrolyze ATP within the timeframe of this experiment (half-life >30min, [28]) and further confirmed by our ATP hydrolysis activity assays (Fig. 3). The TR-SAXS kinetic profile of the purified TM287/288 E/Q-mutant is almost identical to that of the wild-type transporter (Suppl. Fig. S4A) indicating that ATP hydrolysis and resulting conformational changes can be ignored in our further analyses. Thus, our analysis covers the conformational changes from IF (start) to OF (end) without return.

**Figure 3.**
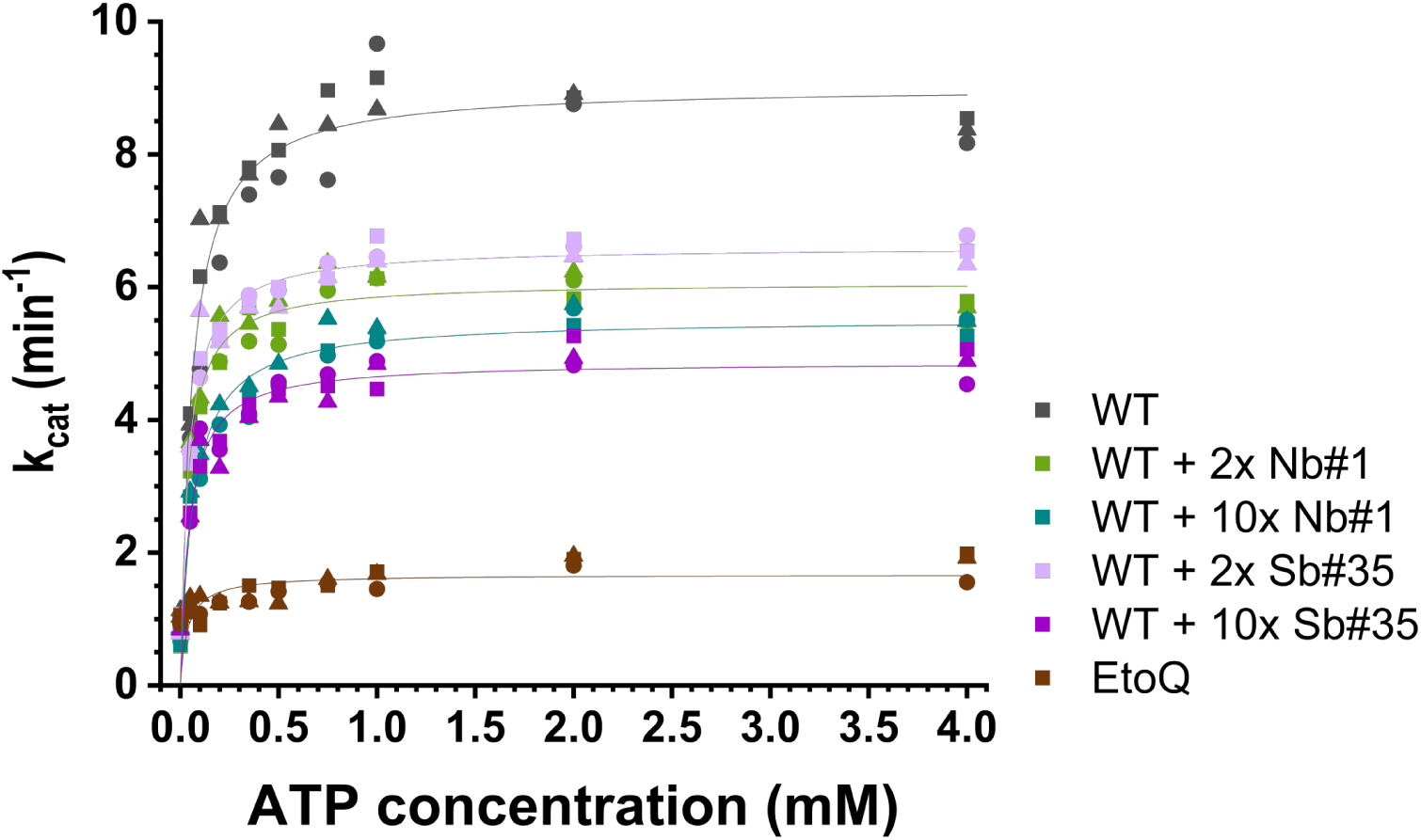
Activity assays and binding data of TM287/288 and conformation-specific single-domain antibodies. Activity assays showing decrease in activity of TM287/288 when bound to conformation specific single-domain antibodies Nb#1 or Sb#35. All assays were performed as triplicates represented by the different symbols for each data point. The data was then fitted to the Michaelis-Menten equation.

This is further supported by the results of our activity assays which show that ATP is hydrolyzed by the wild-type protein at room temperature (Fig. 3). However, hydrolysis is slow at RT (k_cat_ = 9 min^-1^ /0.15 s^-1^) because TM287/288 originates from *T. maritima* which has its temperature optimum around 80°C. Thus, we attribute the second phase of increase in R_g_ beginning around 30 s to the transition between occluded state (Occ) and outward-facing (OF) state of the transporter (Fig. 2B).

The conditions required for an ABC transporter to adopt its outward-facing (OF) conformation — the high-energy state of its transport cycle — remain under debate. Some studies argue that ATP hydrolysis is essential for OF formation [46, 47], while others report that ATP binding alone suffices for the IF-to-OF transition [15–17, 48]. Our SF-TR-SAXS data support the latter, indicating that ATP binding alone drives this conformational change, consistent with the ATP-switch model [10].

In order to quantify the changes in R_g_ magnitude during the TM287/288 conformational cycle, we generated models of TM287/288 in DDM micelles based on available structures, and homology models in various conformational states (Suppl. Fig. S3). We then calculated R_g_ values for these models using CRYSOL [40] (see table in Suppl. Fig. S3), calculated ΔR_g_ values between different states, and compared these Δ R_g_ values to our experimental TR-SAXS data. We compared ΔR_g_ rather than absolute R_g_ values in order to cancel out differences due to different protein constructs (e.g. His-tag and linker). Using the IF – Occ – OF transition of TM287/288 shown in Figure 2 as example, the R_g_ values change from 47.9 Å (IF starting structure) to 44.0 Å (Occ transition state) to 45.4 Å (OF state). The corresponding ΔR_g_ values of -3.9 Å and -2.5 Å agree well with our observed TR-SAXS data (Fig. 2). The observed SAXS trace also supports the assumption of a wider open IF starting state than the current crystal structures of TM287/288 as a more closed IF starting state (such as pdb:4Q4A) would not fit the data well. A closed IF starting state would result in a calculated ΔR_g_ of -0.2 Å and thus not agree with our observed experimental data (Fig. 2/ Suppl. Fig. S3). Our kinetic SAXS data reveal that TM287/288 undergoes a two-step conformational change upon nucleotide addition: a rapid compaction phase leading to a minimal R_g_ around 20–30 seconds, followed by a slower expansion phase that plateaus between 2–4 minutes. These observations offer mechanistic insight into the sequential formation of an occluded intermediate and its transition to the OF state.

### Conformational trapping of TM287/288 using state-specific single-domain antibodies

Recently, two different single-domain antibodies were generated that bind TM287/288 in the presence of ATP and inhibit its ATPase activity. In the crystal structures, both nanobodies bind the transporter in the OF state, with either nanobody Nb#1 bound to the bottom of the closed NBD dimer (pdb:6QV1) or sybody Sb#35 bound to the top of an extracellular wing (pdb:6QUZ) [28]. They showed that Nb#1 and Sb#35 reduce ATPase activity by about 30 % and 70 %, respectively, at 10x molar excess [28] which agrees well with our activity data shown in Fig. 3.

We sought to use these conformation-specific nanobodies as markers of different TM287/288 states in our TR-SAXS experiments. In order to investigate their binding kinetics and the associated conformational changes, we mixed TM287/288 with either Nb#1 or Sb#35 and then added Mg^2+^-ATP via stopped-flow mixing followed by SAXS profile acquisition. In previous studies it was shown that both single domain nanobodies (Nb#1 / Sb#35) bind preferentially to ATP-bound TM287/288 with some weak affinity binding of Nb#1 also to apo TM287/288 at high concentrations [28, 49]. Our static SAXS data, however, indicate no binding of Nb#1 to apo TM287/288 in the absence of ATP (Suppl. Fig. S5), thus we assume that their binding epitopes will only become accessible after ATP-induced conformational changes of the transporter.

For Nb#1, which binds at the interface of the dimerised NBDs, we observed a decrease in R_g_ during the first 20 s, which is attributed to the formation of an occluded state of TM287/288. This step is followed by a rapid increase in R_g_, peaking around 45 s which we associate with the binding of Nb#1 to the closed NBD dimer and a subsequent decrease in R_g_ over 15 s. No further change in R_g_ is observed up to the measured timepoint of 240 s (Fig. 4A).

**Figure 4.**
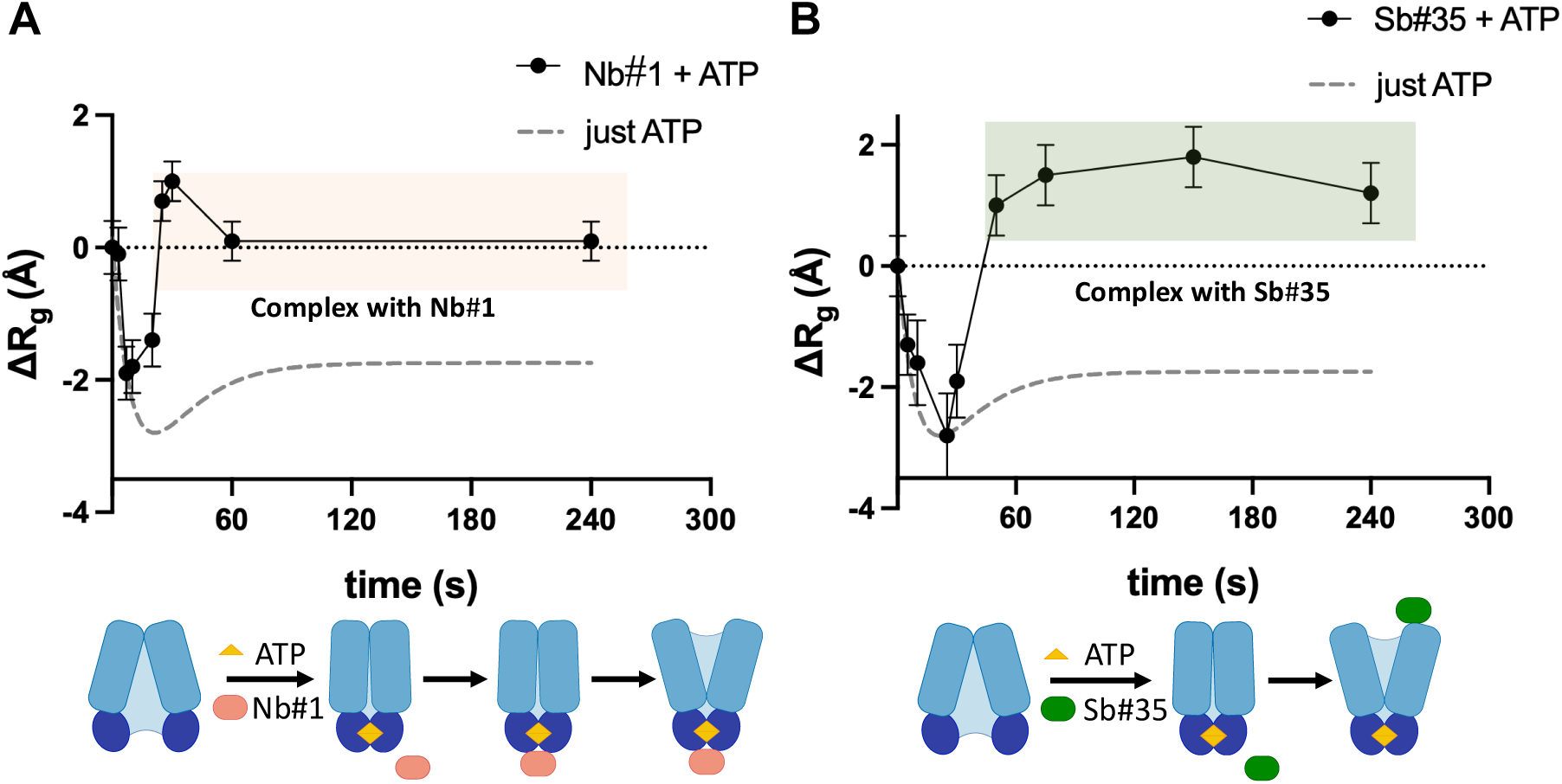
Using single-domain antibodies as conformational markers during TM287/288 turnover. SF-TR-SAXS data acquired for TM287/288 (20 μM) premixed with Nb#1 (A) or Sb#35 (B) (40 μM) before SF-mixed with ATP-Mg^2+^ (1 mM). Nb#1 binds to the NBDs of TM287/288 in the Occ state while Sb#35 binds to the extracellular side only in the OF state.

In the case of Sb#35, our TR-SAXS data show a similar overall trend with initial decrease in the R_g_ of the transporter, followed by subsequent increase in R_g_ but at a later time than with Nb#1 (Fig. 4B). The plateau reached from 80 s shows larger R_g_ values than calculated from the model, most likely caused by minor (radiation induced) aggregation of excess Sb#35.

Comparing the timescales of R_g_ changes in the presence of nanobody/sybody with those of apo TM287/288 with Mg^2+^-ATP alone, it is evident that Nb#1 already binds to the occluded state of TM287/288 (to NBDs), while Sb#35 only binds to the OF state (from extracellular side). This agrees with the fact that the binding site for Nb#1, the closed NBD dimer, is formed in the occluded transporter conformation and is already accessible to the Nb#1. In contrast, the binding site for Sb#35, which is formed by the extracellular wing of TM287/288 only becomes accessible after transition of the transporter from the occluded state to the OF state. This explains the delayed rise in R_g_ from Sb#35 association during our mixing experiments (Suppl. Fig. S6).

The binding kinetics of the nanobodies had already been determined previously by SPR (Sb#35: k_on_ = 1.43*10^4^ M^-1^ s^-1^, k_off_ = 1.57*10^-3^ s^-1^ / Nb#1: k_on_ = 3.41*10^5^ M^-1^ s^-1^, k_off_ = 6.28*10^-2^ s^-1^) [28]. However, binding kinetics of the nanobodies are unlikely to play a major role in this TR-SAXS setup, because the association phase of binding is very fast due to the high Nb/Sb concentrations (20 μM; approx. 1000 x K_d_). The off-rate would only play a role under non-equilibrium situations (i.e. if free nanobodies are removed from the system), which is not the case in our setup.

An unresolved question concerns the “overshoot” around 45 s observed for the TM287/288+Nb#1 complex. This overshoot which is absent in the complex with Sb#35 (Suppl. Fig. S6) could be explained by a 2-step association mechanism of Nb#1 with changes in the NBD dimer interface.

Additionally, we could further show that both single-domain antibodies (Nb#1/Sb#35) can bind to TM287/288 simultaneously after mixing with Mg^2+^-ATP, resulting in a large increase in R_g_, consistent with our model of the ternary state (Suppl. Fig. S7A). The calculated R_g_ of the ternary state (TM287/288+Nb#1+Sb#35) is 48.9 Å (Suppl. Fig. S3). The simultaneous binding of both single-domain antibodies to TM287/288 is further supported by Biolayer Interferometry (BLI) data (Suppl. Fig. S7B). In this experiment we immobilized biotinylated TM287/288 to the BLI sensor and measured binding to Nb#1 in the first step, resulting in a rapid signal increase (large k_on_). In the second step, we further added Sb#35 and observed an additional signal increase (with slower k_on_) indicating simultaneous binding of both single domain antibodies. The differences in association kinetics of Nb#1 and Sb#35 to TM287/288 in the ternary complex agree with those previously observed for the binary complexes (see above) [28].

## Conclusions

Altogether, our SF-TR-SAXS experiments provide a detailed mechanistic insight into the structural kinetics of ATP-driven conformations of the ABC transporter TM287/288. By observing changes in the transporter R_g_ we can track transitions between its catalytic states on the second-to-minute timescale. To validate our findings, we make use of conformation-selective single-domain antibodies as temporal markers of different structural states in solution. With the advent of *de novo* binder designs targeting different protein conformations using machine learning methods, techniques such as TR-SAXS can provide detailed structural kinetic insights into the molecular function of numerous proteins.

Upon rapid mixing with Mg²⁺-ATP, we observed an immediate decrease in Rg, reaching a minimum within ∼20–30 seconds. This compaction is consistent with ATP-driven dimerization of the nucleotide-binding domains (NBDs) and formation of an occluded state, which until now had only been predicted by simulations for TM287/288. Whether this occluded state resembles a fully-occluded state or an outward-facing occluded state as previously shown for other ABC transporters [19, 20, 24] cannot be determined using this setup as their R_g_ values are very similar. Importantly, the subsequent partial re-expansion in R_g_ suggests a transition from the Occ to the OF state. This is in agreement with spontaneous large-scale conformational transitions from IF via Occ to OF conformation that have been observed by MD simulations [45]. This two-step progression, unaffected in a catalytically inactive E/Q mutant, confirms that SF-TR-SAXS can resolve nucleotide-dependent conformational changes independent of hydrolysis within minutes. Comparison with calculated R_g_ values from structural models further supports the assignment of each state.

In addition, we exploited state-specific nanobodies (Nb#1 and Sb#35) as conformational traps. Their distinct binding kinetics and SAXS profiles reinforce the detection of the Occ and OF states. The ability to capture simultaneous binding of both nanobodies highlights the sequential accessibility of these conformations.

Interestingly, the conformation-specific nanobodies do not accelerate the formation of the OF occluded or OF open conformations. This may be explained by the “forward-only” reaction coordinate meaning that once ATP is bound, the reaction cannot reverse. In addition, it proofs that nanobodies/sybodies do not artificially push the target protein into a new conformation, but instead only stabilize a conformation once it is formed (= conformational trapping).

Together, these results demonstrate that SF-TR-SAXS is a powerful tool to resolve short-lived intermediates that are central to the transport mechanism. By combining rapid mixing with high-resolution structural modeling and conformationally selective nanobodies, we establish a robust framework to characterize dynamic states in ABC transporters and other complex membrane proteins.

## Author Contributions

Conceptualization, I.J. and H.T.; Methodology, I.J., C.E.B., and H.T.; Investigation, L.S., A.G., L.V.S., D.D., C.E.B., M.A.S., H.T and I.J..; Writing – Original Draft, I.J. and H.T.; Writing – Review & Editing, all authors; Funding Acquisition, L.V. S., M.A.S., and H.T.; Supervision, H.T. and I.J.

## Acknowledgements

We are grateful to the staff at the beamline P12 (EMBL, Hamburg) and acknowledge access to the Sample Preparation and Characterization (SPC) Facility of EMBL, Hamburg. We thank Melanie Scherer and Cedric Hutter for plasmids and protocols. This research was funded by a Heisenberg grant (to HT) and the Cluster of Excellence ‘Advanced Imaging of Matter’ of the Deutsche Forschungsgemeinschaft (DFG) - EXC 2056 - project ID 390715994.

## Footnotes

ABC: ATP-binding cassette
ATP: adenosine triphosphate
DDM: dodecylmaltoside
IMAC: immobilized metal affinity chromatography
IMP: integral membrane protein
MD: molecular dynamics
Nb: nanobody
NBD: nucleotide-binding domain
R_g_: radius of gyration
SAXS: small-angle X-ray scattering
Sb: sybody
SF: stopped-flow
TMD: transmembrane domain
TR: time-resolved

## References

1. Alam, A. and K.P. Locher, Structure and Mechanism of Human ABC Transporters. Annu Rev Biophys, 2023.

2. Dean, M., A. Rzhetsky, and R. Allikmets, The human ATP-binding cassette (ABC) transporter superfamily. Genome Res, 2001. 11(7): p. 1156–66.

3. Locher, K.P., Mechanistic diversity in ATP-binding cassette (ABC) transporters. Nat Struct Mol Biol, 2016. 23(6): p. 487–93.

4. Cui, J. and A.L. Davidson, ABC solute importers in bacteria. Essays Biochem, 2011. 50(1): p. 85–99.

5. Davidson, A.L., et al., Structure, function, and evolution of bacterial ATP-binding cassette systems. Microbiol Mol Biol Rev, 2008. 72(2): p. 317–64, table of contents.

6. Higgins, C.F., The ABC of channel regulation. Cell, 1995. 82(5): p. 693–6.

7. Schmitt, L. and R. Tampe, Structure and mechanism of ABC transporters. Curr Opin Struct Biol, 2002. 12(6): p. 754–60.

8. Thomas, C. and R. Tampe, Structural and Mechanistic Principles of ABC Transporters. Annu Rev Biochem, 2020. 89: p. 605–636.

9. Rees, D.C., E. Johnson, and O. Lewinson, ABC transporters: the power to change. Nat Rev Mol Cell Biol, 2009. 10(3): p. 218–27.

10. Higgins, C.F. and K.J. Linton, The ATP switch model for ABC transporters. Nat Struct Mol Biol, 2004. 11(10): p. 918–26.

11. Janas, E., et al., The ATP hydrolysis cycle of the nucleotide-binding domain of the mitochondrial ATP-binding cassette transporter Mdl1p. J Biol Chem, 2003. 278(29): p. 26862–9.

12. Jardetzky, O., Simple allosteric model for membrane pumps. Nature, 1966. 211(5052): p. 969–70.

13. Lewinson, O., C. Orelle, and M.A. Seeger, Structures of ABC transporters: handle with care. FEBS Lett, 2020. 594(23): p. 3799–3814.

14. Orelle, C., L. Schmitt, and J.M. Jault, Waste or die: The price to pay to stay alive. Trends Microbiol, 2023. 31(3): p. 233–241.

15. Hofmann, S., et al., Conformation space of a heterodimeric ABC exporter under turnover conditions. Nature, 2019. 571(7766): p. 580–583.

16. Nocker, C., et al., Single-molecule dynamics reveal ATP binding alone powers substrate translocation by an ABC transporter. bioRxiv, 2025.

17. Stefan, E., S. Hofmann, and R. Tampe, A single power stroke by ATP binding drives substrate translocation in a heterodimeric ABC transporter. Elife, 2020. 9.

18. Kehlenbeck, D.M., et al., Cryo-EM structure of MsbA in saposin-lipid nanoparticles (Salipro) provides insights into nucleotide coordination. FEBS J, 2022. 289(10): p. 2959–2970.

19. Li, P., et al., Structures of Atm1 provide insight into [2Fe-2S] cluster export from mitochondria. Nat Commun, 2022. 13(1): p. 4339.

20. Lin, D.Y., S. Huang, and J. Chen, Crystal structures of a polypeptide processing and secretion transporter. Nature, 2015. 523(7561): p. 425–30.

21. Mi, W., et al., Structural basis of MsbA-mediated lipopolysaccharide transport. Nature, 2017. 549(7671): p. 233–237.

22. Thomas, C. and R. Tampe, Multifaceted structures and mechanisms of ABC transport systems in health and disease. Curr Opin Struct Biol, 2018. 51: p. 116–128.

23. Carrillo, V.H.P., et al., Bidirectional communication between nucleotide and substrate binding sites in a type IV multidrug ABC transporter. Nat Commun, 2025. 16(1): p. 9921.

24. Stefan, E., et al., De novo macrocyclic peptides dissect energy coupling of a heterodimeric ABC transporter by multimode allosteric inhibition. Elife, 2021. 10.

25. Procko, E., et al., The mechanism of ABC transporters: general lessons from structural and functional studies of an antigenic peptide transporter. FASEB J, 2009. 23(5): p. 1287–302.

26. Hohl, M., et al., Crystal structure of a heterodimeric ABC transporter in its inward-facing conformation. Nat Struct Mol Biol, 2012. 19(4): p. 395–402.

27. Hohl, M., et al., Structural basis for allosteric cross-talk between the asymmetric nucleotide binding sites of a heterodimeric ABC exporter. Proc Natl Acad Sci U S A, 2014. 111(30): p. 11025–30.

28. Hutter, C.A.J., et al., The extracellular gate shapes the energy profile of an ABC exporter. Nat Commun, 2019. 10(1): p. 2260.

29. Levantino, M., et al., Using synchrotrons and XFELs for time-resolved X-ray crystallography and solution scattering experiments on biomolecules. Curr Opin Struct Biol, 2015. 35: p. 41–8.

30. Akiyama, S., et al., Conformational landscape of cytochrome c folding studied by microsecond-resolved small-angle x-ray scattering. Proc Natl Acad Sci U S A, 2002. 99(3): p. 1329–34.

31. Arai, M., et al., Microsecond hydrophobic collapse in the folding of Escherichia coli dihydrofolate reductase, an alpha/beta-type protein. J Mol Biol, 2007. 368(1): p. 219–29.

32. Konuma, T., et al., Time-resolved small-angle X-ray scattering study of the folding dynamics of barnase. J Mol Biol, 2011. 405(5): p. 1284–94.

33. West, J.M., et al., Time evolution of the quaternary structure of Escherichia coli aspartate transcarbamoylase upon reaction with the natural substrates and a slow, tight-binding inhibitor. J Mol Biol, 2008. 384(1): p. 206–18.

34. Josts, I., et al., Structural Kinetics of MsbA Investigated by Stopped-Flow Time-Resolved Small-Angle X-Ray Scattering. Structure, 2020. 28(3): p. 348–354 e3.

35. Blanchet, C.E., et al., Versatile sample environments and automation for biological solution X-ray scattering experiments at the P12 beamline (PETRA III, DESY). J Appl Crystallogr, 2015. 48(Pt 2): p. 431–443.

36. Franke, D., C.M. Jeffries, and D.I. Svergun, Correlation Map, a goodness-of-fit test for one-dimensional X-ray scattering spectra. Nat Methods, 2015. 12(5): p. 419–22.

37. Franke, D., et al., ATSAS 2.8: a comprehensive data analysis suite for small-angle scattering from macromolecular solutions. J Appl Crystallogr, 2017. 50(Pt 4): p. 1212–1225.

38. Webb, B. and A. Sali, Protein Structure Modeling with MODELLER. Methods Mol Biol, 2021. 2199: p. 239–255.

39. Cheng, X., et al., CHARMM-GUI micelle builder for pure/mixed micelle and protein/micelle complex systems. J Chem Inf Model, 2013. 53(8): p. 2171–80.

40. Svergun, D.I., C. Barberato, and M.H.J. Koch, CRYSOL - a Program to Evaluate X-ray Solution Scattering of Biological Macromolecules from Atomic Coordinates. J. Appl. Crystallogr., 1995. 28: p. 768–773.

41. Baginski, E.S., E. Epstein, and B. Zak, Review of phosphate methodologies. Ann Clin Lab Sci, 1975. 5(5): p. 399–416.

42. Chifflet, S., et al., A method for the determination of inorganic phosphate in the presence of labile organic phosphate and high concentrations of protein: application to lens ATPases. Anal Biochem, 1988. 168(1): p. 1–4.

43. Kehlenbeck, D.M., et al., Comparison of lipidic carrier systems for integral membrane proteins - MsbA as case study. Biol Chem, 2019. 400(11): p. 1509–1518.

44. Goddeke, H. and L.V. Schafer, Capturing Substrate Translocation in an ABC Exporter at the Atomic Level. J Am Chem Soc, 2020. 142(29): p. 12791–12801.

45. Goddeke, H., et al., Atomistic Mechanism of Large-Scale Conformational Transition in a Heterodimeric ABC Exporter. J Am Chem Soc, 2018. 140(13): p. 4543–4551.

46. Mishra, S., et al., Conformational dynamics of the nucleotide binding domains and the power stroke of a heterodimeric ABC transporter. Elife, 2014. 3: p. e02740.

47. Verhalen, B., et al., Energy transduction and alternating access of the mammalian ABC transporter P-glycoprotein. Nature, 2017. 543(7647): p. 738–741.

48. Timachi, M.H., et al., Exploring conformational equilibria of a heterodimeric ABC transporter. Elife, 2017. 6.

49. Galazzo, L., et al., Spin-labeled nanobodies as protein conformational reporters for electron paramagnetic resonance in cellular membranes. Proc Natl Acad Sci U S A, 2020. 117(5): p. 2441–2448.

