## Supplementary Information for "Capturing transient states of heterodimeric ABC transporter TM287/288 by Time-Resolved Small-Angle X-ray Scattering"

### Supplementary figures

### <u>Suppl. Figure S1</u>

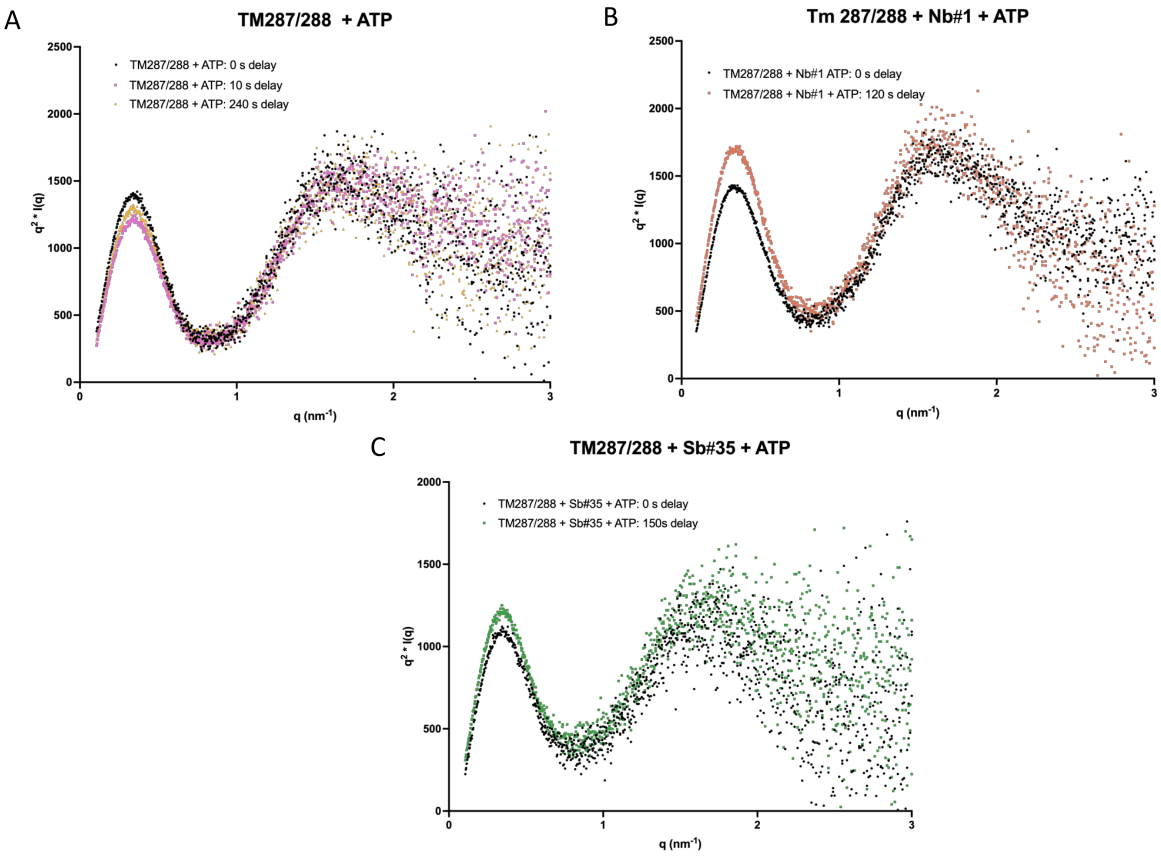


**Kratky plots of representative experimental SAXS curves at selected time points.** A) Three time points of TM287/288 binding ATP-$Mg^{2+}$ representing evolution of conformational changes in the sample. B) Two time points showing changes in scattering of TM287/288 upon binding ATP and Nb#1. C) Two time points showing changes in scattering of TM287/288 upon binding ATP and Sb#35.

**Suppl. Figure S2**

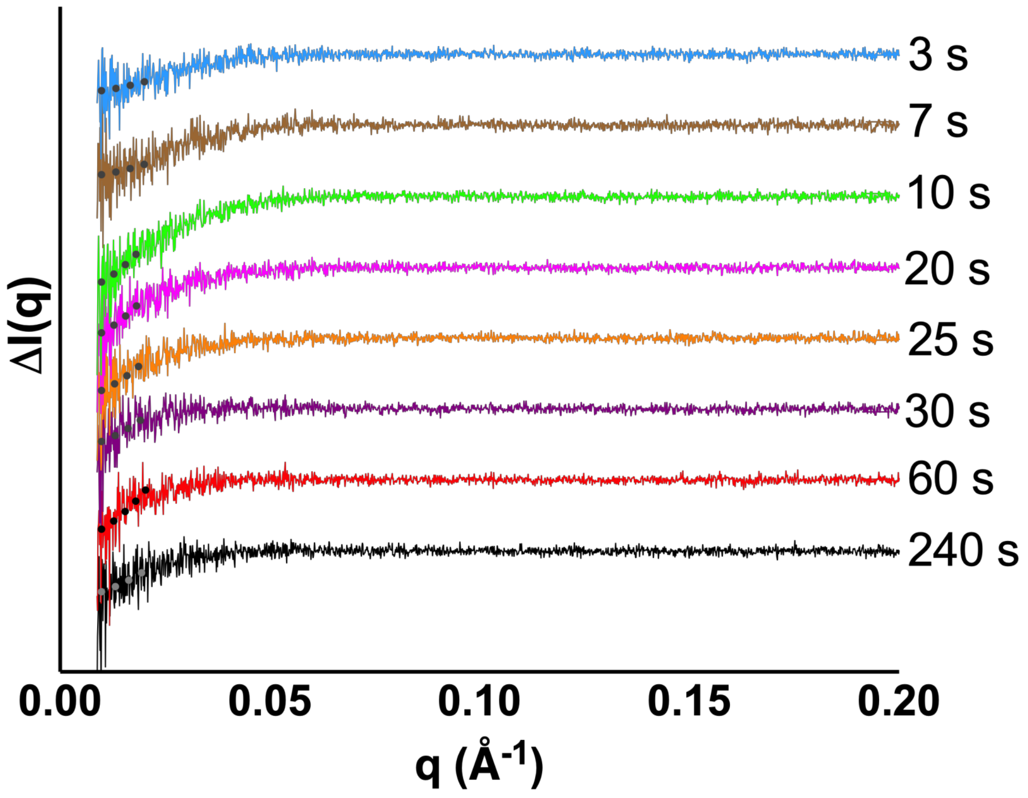


**Time-resolved scattering difference curves show conformational changes in TM287/288 upon ATP binding.** Stacked Δ I(q) curves showing changes in scattering signal over reaction time. Dotted lines emphasise changes in low-q region where Guinier fits were calculated.

### Suppl. Figure S3

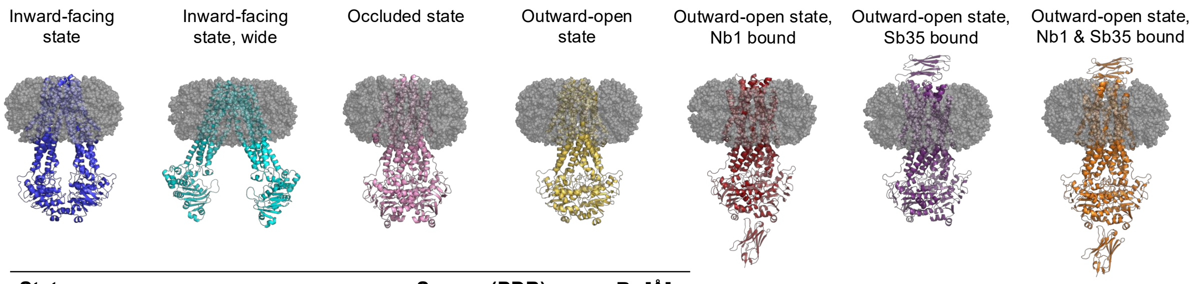


| State | Source (PDB) | $R_g$ [Å] |
|---|---|---|
| Inward-facing | 4Q4A | 44.2 |
| Inward-facing, wide | 6BL6 | 47.9 |
| Occluded | modelled from: 4S0F, 7PR1, 6RAI | 44.0 |
| Outward-open | 6QUZ | 45.4 |
| Outward-open, Nb1 bound | modelled from: 6QV1 | 48.5 |
| Outward-open, Sb35 bound | modelled from: 6QUZ | 46.0 |
| Outward-open, Nb1 & Sb35 bound | modelled from: 6QV1, 6QUZ | 48.9 |

**Illustration of different conformational states of TM287/288 in detergent (DDM) micelles with $R_g$ values indicated.** PDB codes of structures or templates for models are given in the table. inward-facing state, blue / wide inward-facing apo state, cyan / occluded state, pink / outward-open state, yellow / outward-open, Nb1 bound state, red / outward-open Sb35-bond state, violet / outward-open Nb1 and Sb35-bound state, orange. Detergent micelles were added using CHARMM-GUI micelle builder (Cheng, Jo et al. 2013), and $R_g$ values were calculated using CRYSOL (Svergun, Barberato et al. 1995).

**Suppl. Figure S4**

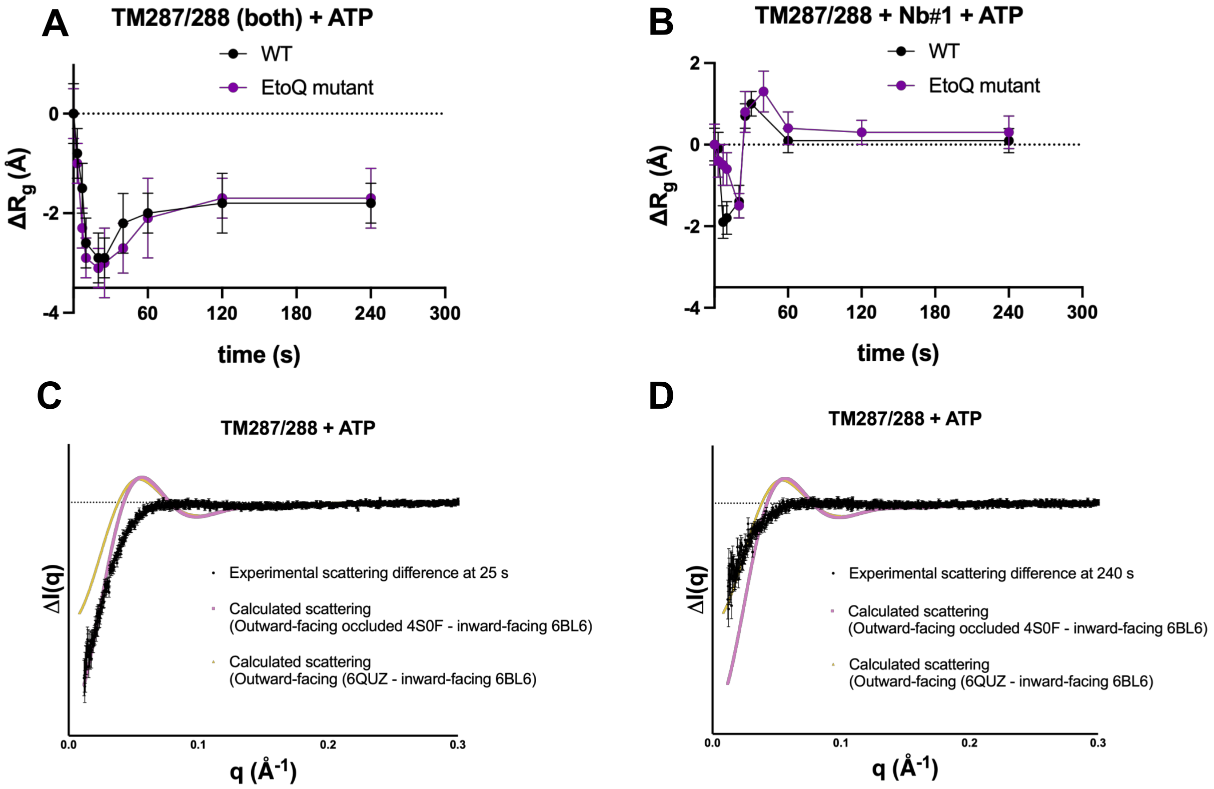


**SF-TR-SAXS comparison of wt TM287/288 and its E/Q-mutant (E517Q).** A) Apo proteins mixed with ATP-$Mg^{2+}$. B) TM287/288 variants premixed with Nb#1 before SF-mixed with ATP-$Mg^{2+}$. C) and D) Comparison of the 25 s time point (where the occluded state is thought to populate) and 240 s time point (where outward-open state is thought to populate) in our kinetic reaction with two calculated models of TM287/288. The $\chi^2$ values for the fit between 25 s and Occ model is 13.8 versus 26.7 for OF model. At 240 s the $\chi^2$ for Occ is 10 but 3.9 for OF model. The 240 s difference dataset is noisier and larger experimental uncertainties lead to lower $\chi^2$ values.

**Suppl. Figure S5**

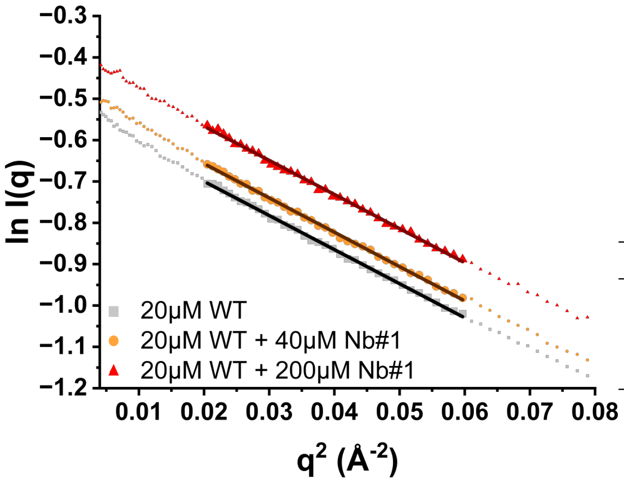


| sample | slope Guinier plot | $R_g$ [Å] |
|---|---|---|
| TM287/288 WT | -8.25 ± 0.05 | 49.7 ± 0.2 |
| TM287/288 WT + 2x Nb#1 | -8.30 ± 0.05 | 49.9 ± 0.2 |
| TM287/288 WT + 10x Nb#1 | -8.25 ± 0.07 | 49.7 ± 0.3 |

**Guinier analysis of apo TM287/288 in absence and presence of Nb#1 indicate that Nb#1 is not binding to TM287/288 in the absence of $Mg^{2+}$-ATP.**

**Suppl. Figure S6**

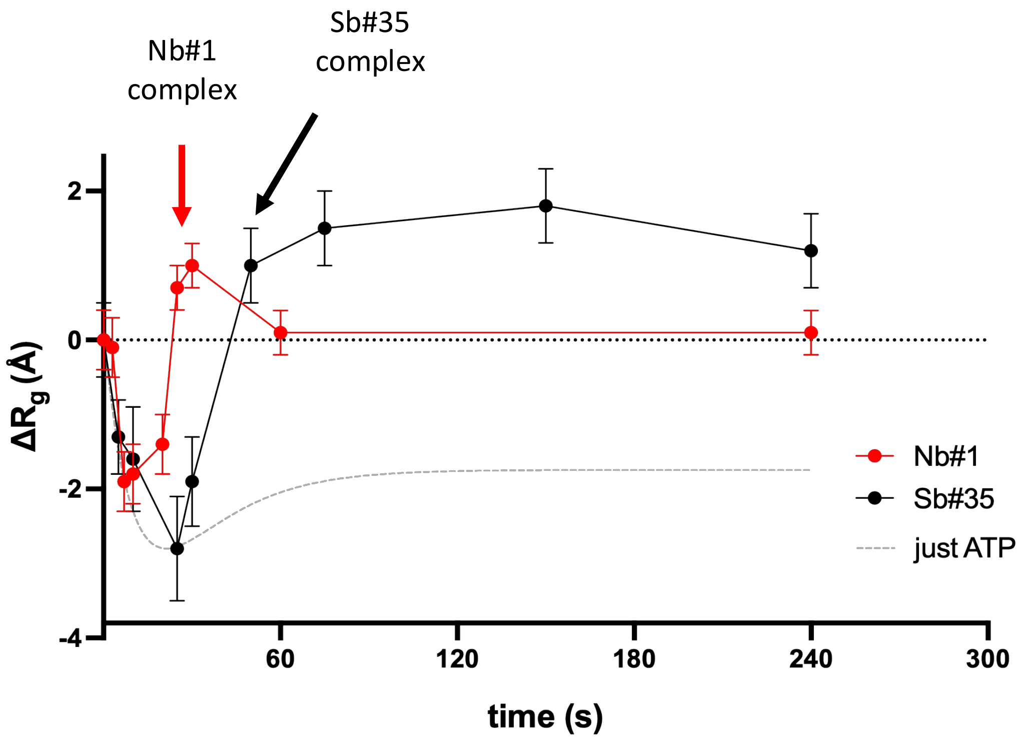


**SF-TR-SAXS comparison of TM287/288 in complex with Nb#1 and Sb#35.** SF-TR-SAXS data acquired for TM287/288 premixed with Nb#1 (A) or Sb#35 (B) before SF-mixed with ATP-$Mg^{2+}$. Nb#1 binds to the NBDs of TM287/288 in the occluded state while Sb#35 binds to the extracellular side only in the OF state.

**Suppl. Figure S7**

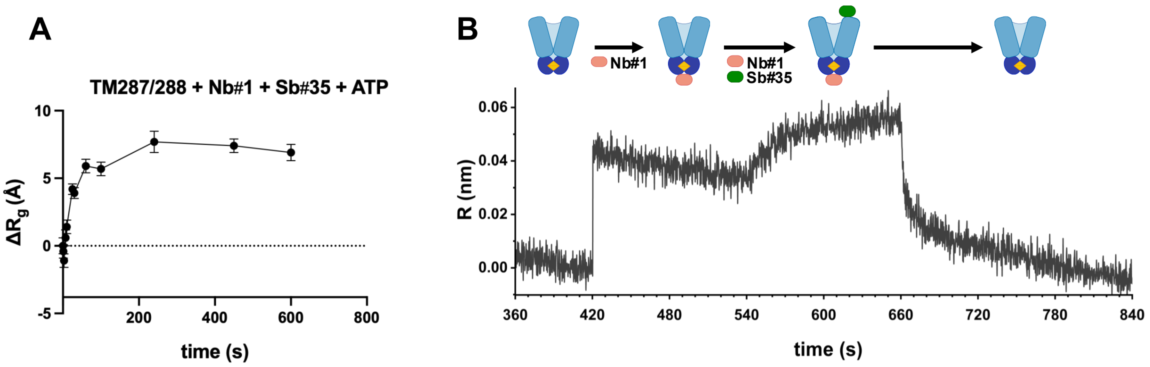


**Simultaneous binding of both single-domain antibodies to TM287/288.** A) SF-TR-SAXS data acquired for TM287/288 premixed with both Nb#1 and Sb#35 before SF-mixed with ATP-$Mg^{2+}$. B) BLI measurements showing association of Nb#1 and Sb#35 followed by their dissociation.

**Suppl. Table 1**

| | $K_m$ | $v_{max}$ | $k_{cat}$ |
|---|---|---|---|
| | *mM* | *$nmol_{ATP}/min/mg_{TM287/288}$* | *$min^{-1}$* |
| **wt** | 0.059 ± 0.008 | 67.7 ± 1.5 | 9.02 ± 0.20 |
| **wt 2x Nb1** | 0.039 ± 0.005 | 45.6 ± 0.8 | 6.07 ± 0.11 |
| **wt 10x Nb1** | 0.066 ± 0.008 | 41.4 ± 0.9 | 5.52 ± 0.12 |
| **wt 2x Sb35** | 0.041 ± 0.005 | 49.5 ± 0.9 | 6.61 ± 0.12 |
| **wt 10x Sb35** | 0.048 ± 0.008 | 36.5 ± 0.9 | 4.87 ± 0.12 |
| **E517Q** | 0.038 ± 0.018 | 12.5 ± 0.8 | 1.67 ± 0.11 |

**Suppl. Table S1: ATPase activity assays.** Activity assays for TM287/288 were performed using the Baginski method at room temperature (Baginski, Epstein et al. 1975, Chifflet, Torriglia et al. 1988). $K_m$ and $k_{cat}$ were calculated from fitting the experimental data according to Michaelis-Menten. Standard errors were obtained from triplicate measurements.

### References


Baginski, E. S., E. Epstein and B. Zak (1975). "Review of phosphate methodologies." Ann Clin Lab Sci **5**(5): 399-416.

Cheng, X., S. Jo, H. S. Lee, J. B. Klauda and W. Im (2013). "CHARMM-GUI micelle builder for pure/mixed micelle and protein/micelle complex systems." J Chem Inf Model **53**(8): 2171-2180.

Chifflet, S., A. Torriglia, R. Chiesa and S. Tolosa (1988). "A method for the determination of inorganic phosphate in the presence of labile organic phosphate and high concentrations of protein: application to lens ATPases." Anal Biochem **168**(1): 1-4.

Svergun, D. I., C. Barberato and M. H. J. Koch (1995). "CRYSOL - a Program to Evaluate X-ray Solution Scattering of Biological Macromolecules from Atomic Coordinates." J. Appl. Crystallogr. **28**: 768-773.
